# Caffeine increases false alarms in consolidation of face recognition memory

**DOI:** 10.1101/2025.03.27.645742

**Authors:** Candela S. Leon, Agustina L. Lo Celso, Rocío Guajardo, Cecilia Forcato

## Abstract

Numerous studies have shown that caffeine facilitates cognition, particularly memory, when consumed before learning and immediately tested. However, most of this evidence relates to its effects during encoding, and the role in memory consolidation remains unclear. A key study demonstrated that caffeine administered after learning can enhance object recognition memory consolidation by improving the discrimination between previously seen targets and similar lures. Here, we investigated the effects of post-encoding caffeine on the consolidation of face recognition memory using a randomized, double-blind design. Participants (*N* = 104) viewed ten artificially generated faces on Day 1 and then received 200 mg of caffeine or placebo. On Day 2, they completed a recognition task under two conditions: Present condition (original face with five similar distractors) and Absent condition (six similar distractors) adding a “none of the above” option. The results showed that, contrary to our expectation, caffeine consumption did not improve the consolidation of face recognition memory. Instead, it increased False alarm rate in both conditions, reducing the ability to discriminate between previously seen and similar faces. These findings discuss the idea of caffeine as a general cognition enhancer and aligned with studies suggesting it enhances global processing at the expense of detailed discrimination.

## Introduction

Caffeine is the world’s most commonly used psychoactive [1] and nootropic substance [2]. Numerous studies have observed its stimulating effects, which enhance alertness, wakefulness, motivation, motor activity [3,4] and visual attention [5,6] Additionally, caffeine has been shown to mitigate habituation effects [7] and reduce processing speed in inhibitory control tasks [8].

Generally, these studies have focused on caffeine administration before information encoding. As a result, they have not sufficiently distinguished whether its possible enhancing effects influence learning itself or the subsequent consolidation of memories.

Research on caffeine’s effect on memory consolidation, however, remains debated. On the one hand, studies in rodents have demonstrated a positive effect of caffeine when administered post learning [9,10]. Angelucci et al. (2002) [11] showed that relatively low doses (0.3-3 mg/kg) of caffeine improved consolidation in tasks such as the water maze, active avoidance, and inhibitory avoidance. More recently, Días et al. (2022) [12] reported that caffeine enhanced the consolidation of the temporal (’what-when’) and spatial (’what-where’) components of episodic-like memory in rodents. On the other hand, evidence from human studies on the effects of caffeine on memory consolidation remains mixed. One study found that caffeine improved object recognition memory by enhancing the ability to discriminate between familiar and similar objects when the target was absent [13]. In contrast, no improvements were observed in motor memory consolidation following post-practice caffeine administration [14], nor in incidental word learning [15]. Beyond human models, research in invertebrates has also provided nuanced findings. In honey bees, caffeine was found to modulate performance during encoding, with high doses reducing responsiveness, while having no effect on early long-term memory consolidation [16].

While much of the research on caffeine and memory has focused on general or object-based memory tasks, less is known about its effects on more complex and socially relevant forms of memory, such as face recognition. The ability to recognize faces is a widely utilized skill in everyday life. It has been observed that this ability could be modulated by individual variables, such as visual imagery capacity [17]. Furthermore, it could be totally or partially affected in people who acquire or genetically have prosopagnosia (facial blindness) or people with autistic traits [18] The face recognition ability could also be decisive in specific areas such as eyewitness memory, where victims and/or witnesses could play a crucial role in legal decisions through their choices in identification lineups [19] or in the construction of identikits [20]. For this reason, multiple studies have focused on the effect that different legal and illegal drugs have on this ability [21].

To our knowledge, no studies have assessed the influence of caffeine on the consolidation of face recognition memory. Following the study of Borota et al., (2014) [13], we hypothesized that a dose of 200 mg would positively affect human memory consolidation in a recognition task. Therefore, the present study evaluated the effect of caffeine on the consolidation of human face recognition using a randomized double-blind, placebo-controlled study. The experiment was conducted over two days. On Day 1, participants viewed ten artificially generated faces and immediately received either 200 mg of caffeine, the lowest dose that has been proposed to improve consolidation [13], or a placebo. On Day 2, they completed a recognition task under either target present or target absent condition.

## Results

To examine the effects of post-encoding caffeine ingestion on the consolidation of face recognition memory, a randomized, double-blind study was conducted. A total of 104 participants viewed ten artificially generated faces on Day 1 before receiving either 200 mg of caffeine or a placebo. On Day 2, they underwent a recognition task with two conditions: Present (the original face alongside five similar distractors) and Absent (six similar distractors), including a “none of the above” option (Figure 1).

**Figure 1.**
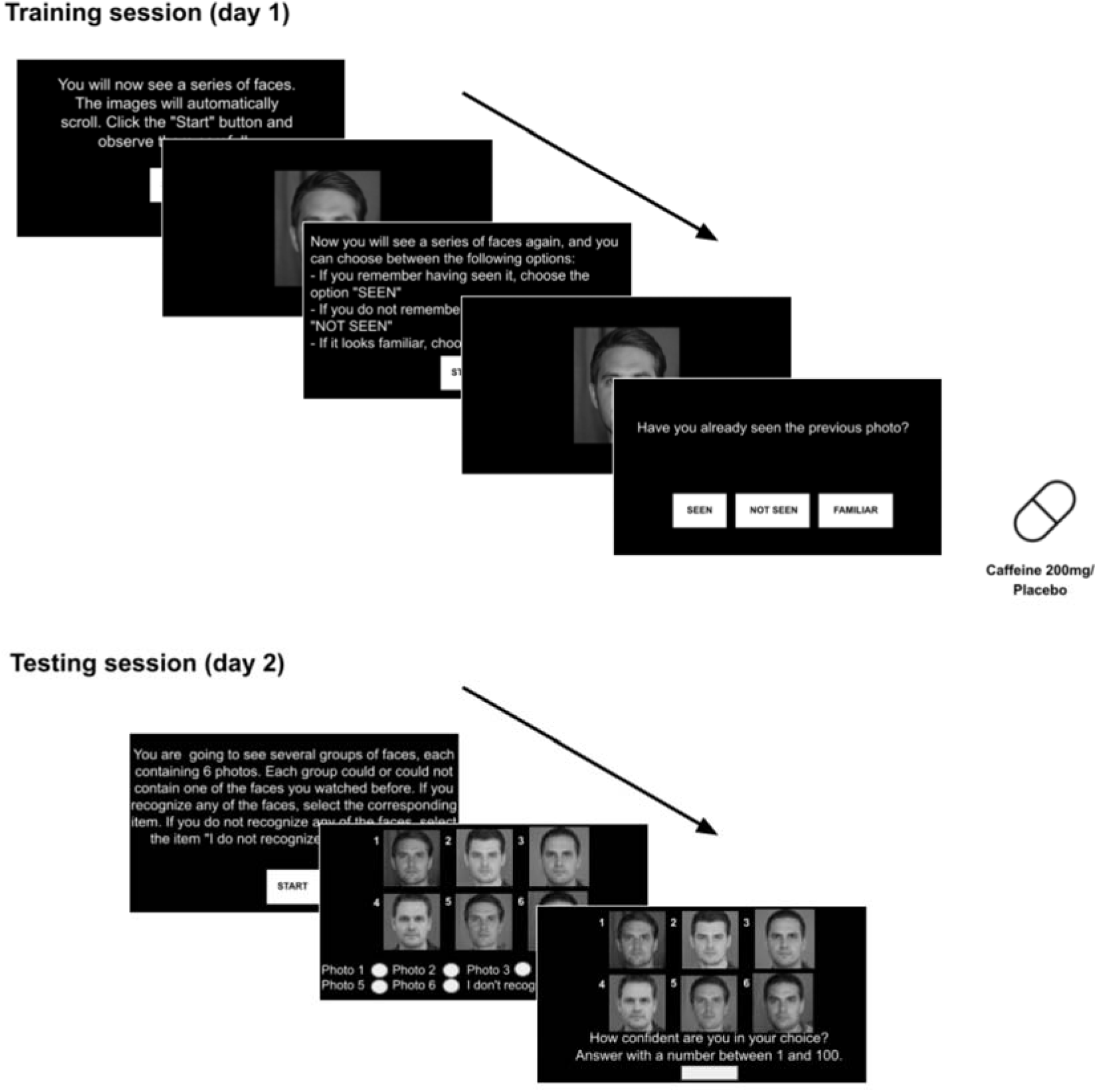
Experimental procedure. On Day 1 participants viewed ten artificially generated faces and immediately received either 200 mg of caffeine or a placebo. On Day 2, they performed a recognition task with two possible conditions: Present (the original face alongside five similar distractors) and Absent (six similar distractors), including a “none of the above” option.

Primarily, the choices made in the Present condition were categorized into hits, false alarms, and incorrect rejections, while in the Absent condition, they were categorized into false alarms and correct rejections for both groups (Table 1).

**Table 1.** Performance Summary in the Testing Session. Percentage of identification outcomes for the Caffeine and Placebo groups on the Testing session (Day 2). Each participant performs 10 identifications.

| Present condition |  |  |  |
| --- | --- | --- | --- |
| Group | Hits | False alarms | Incorrect rejections |
| CAFFEINE | 50.71 % (*N* = 142) | 32.14 % (*N* = 90) | 17.14 % (*N* = 48) |
| PLACEBO | 59.13 % (*N* = 136) | 20.00 % (*N* = 46) | 21.30 % (*N* = 49) |
| Absent condition |  |  |  |
| Group | False alarms | Correct rejections |  |
| CAFFEINE | 70.00 % (*N* = 161) | 29.57 % (*N* = 68) |  |
| PLACEBO | 62.08 % (*N* = 149) | 37.92 % (*N* = 91) |  |

First, sensitivity, i.e. the relationship between correct choices and false alarms, was analyzed for each group in each condition. To analyze the recognition performance in the Present condition, we calculated *d’* value as Z (Hit Rate) – Z (False Alarm Rate). No significant differences were observed between groups (Caffeine group: 0.74 ± 1.34, Placebo group: 1.45 ± 1.51, *t* (48) = −1.75, *p* = 0.08. Fig. 2). Additionally, a one-sample *t* test was conducted to compare the group’s performance against a chance level (*d’* = 0). The results indicated that both groups performed significantly above chance (Caffeine group: 0.74 ± 1.34, *t* (27) = 2.93, *p* = 0.007, 7 0.24, η2 = 0.24. Placebo group: 1.45 ± 1.51, *t* (21) = 4.50, *p* < 0.001, η2 = 0.49. Fig. 2).

**Figure 2.**
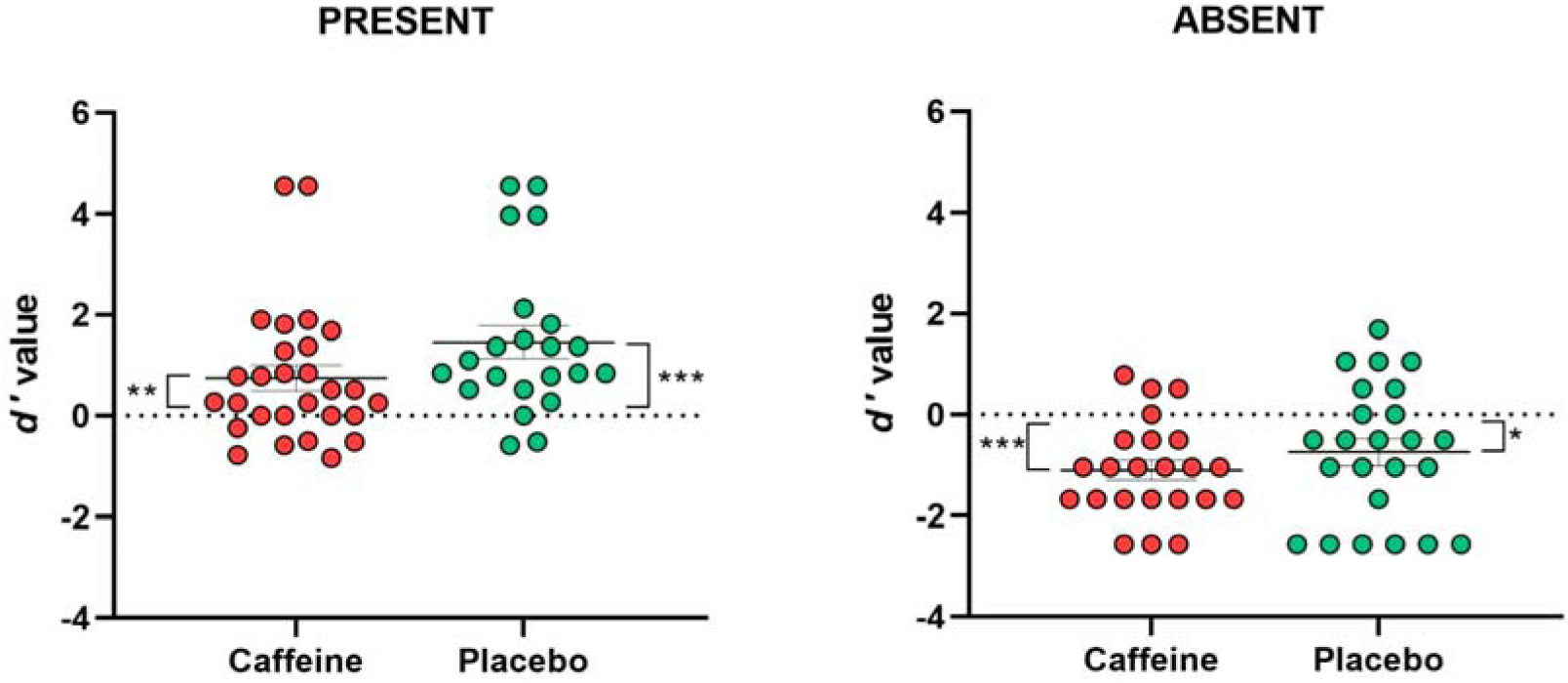
Sensitivity (*d’*) for distinguishing the target (present or remembered) from the fillers across groups for Present and Absent conditions ± SEM. *, *p*□< 0.05. **, *p* < 0.01. ***, *p*□< 0.001.

To analyze the recognition performance in the Absent condition, we calculated *d’* value as Z (Correct rejections) – Z (False Alarm Rate). No significant differences were observed between the groups (Caffeine group: −1.10 ± 0.94, Placebo group: −0.74 ± 1.33, *t* (45) = −1.06, *p* = 0.29, Fig. 2). Furthermore, we found that both groups performed significantly below chance level (*d’* = 0) (Caffeine group: −1.10 ± 0.94, *t* (22) = −5.60, *p* < 0.000. η2 = 0.58. Placebo group: −0.74 ± 1.33, *t* (23) = −2.74, *p* = 0.01. η2 = 0.24. Fig. 2).

To evaluate overall performance across both conditions, we conducted an integrated analysis using a two-way ANOVA with conditions (Present vs. Absent) and group (Caffeine vs. Placebo) as between-subject factors. The dependent variable was the Correct choice rate, defined as the proportion of hits in the Present condition and correct rejections in the Absent condition. This analysis revealed no significant interaction between condition and group (F_condition*group_ (1,89) = 0.08 *p* = 0.76, Fig. 3), indicating that the effect of condition on performance did not differ by group. However, we found that subjects performed better in the Present condition (F_condition_ (1,89) = 30.00 *p* < 0.001, η2 = 0.25, Fig. 3). Furthermore, the Caffeine group showed significantly less correct choices than the Placebo group (F_group_ (1,89) = 4.30 *p* = 0.04, η2 = 0.04, Fig. 3). In the same way, analysis of the False alarm rate revealed no significant interaction between condition and group (F_condition*group_ (1,89) = 0.16 *p* = 0.68, Fig. 3). However, the Absent condition showed a significantly higher False alarm rate compared to the Present condition (F_condition_ (1,89) = 123.13 *p* < 0.001, η2 = 0.57, Fig. 3). Furthermore, the Caffeine group exhibited a higher False alarm rate with respect to Placebo group (F_group_ (1,89) = 7.52 *p* = 0.007, η2 = 0.07, Fig. 3). The variable of Incorrect rejection rate (when in the presence of the original face, they responded that it was none of the options) was only analyzed in the Present condition because the Absent condition does not have this option. It was observed that the groups had no differences for these elections (F_group_ (1,51) = 1.56 *p* = 0.21).

**Figure 3.**
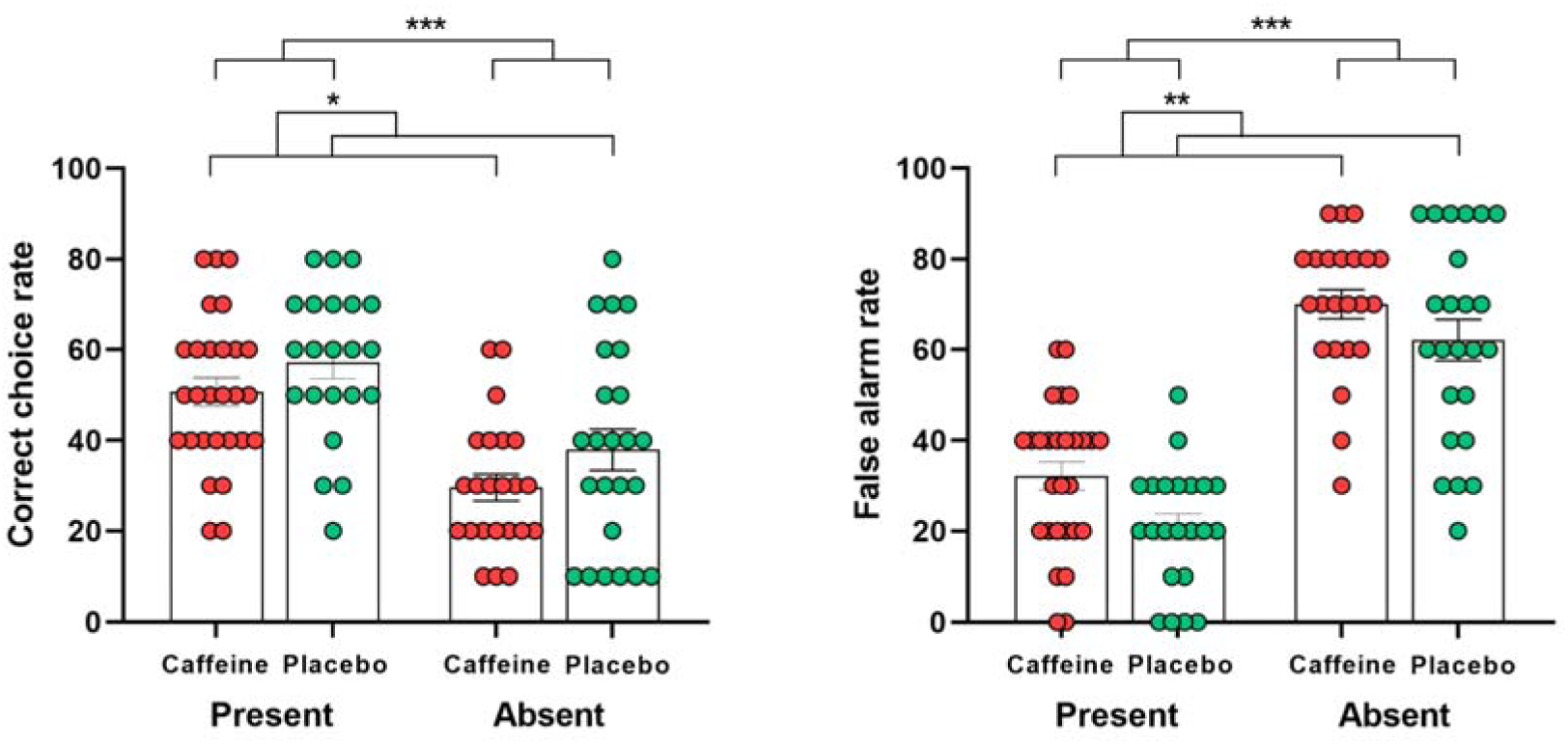
Overall performance across groups and conditions. Correct choice rate (hits and correct rejections) and False alarm rate for both groups across conditions ± SEM. *, *p*□< 0.05. **, *p* < 0.01. ***, *p*□< 0.001

Finally, to ensure that these effects were not influenced by potential confounders, we conducted an additional ANCOVA. On the Correct choice rate, the Covariate BMI (Body mass index) did not have a significant effect (F (1,89) < 0.000, *p* = 0.98), neither state anxiety (State anxiety subscale of the STAI anxiety test) (F (1,89) = 0.33, *p* = 0.56), time awake (Total waking hours from wake-up to the time of the Training session) (F (1,89) = 1.42, *p* = 0.23), or regular daily caffeine consumption (F (1,89) = 2.04, p = 0.15). Further, the same analysis was conducted for the False alarm rate. The Covariate BMI did not have a significant effect (F (1,89) = 0.21, *p* = 0.64), neither state anxiety (F (1,89) = 0.54, *p* = 0.46), time awake (F (1,89) = 1.18, *p* = 0.28), or regular daily caffeine consumption (F (1,89) = 1.65, *p* = 0.20).

Finally, confidence-accuracy characteristic analysis (CAC curves) revealed that the exponential relationship typically observed between responses and confidence is preserved in the Caffeine group (i.e., response rate increases as self-reported confidence increases), despite this group’s lower proportion of correct recognition compared to the placebo group at all confidence levels (Fig. 4).

**Figure 4.**
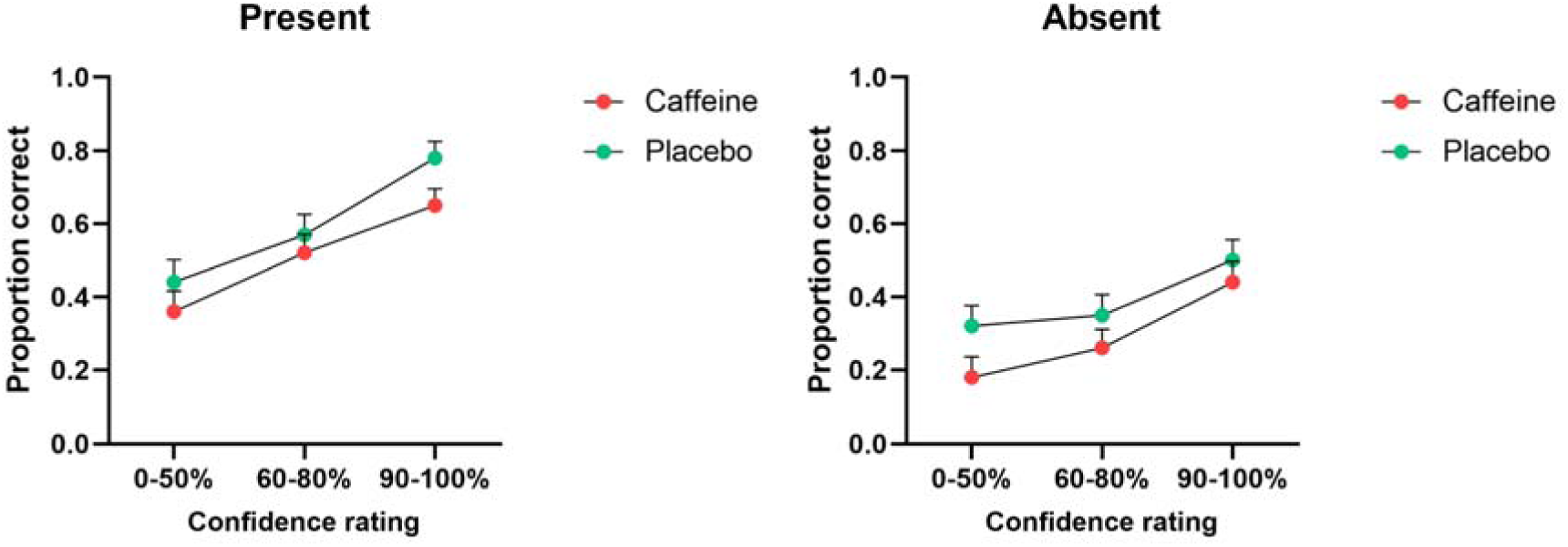
CAC curves for Caffeine and Placebo groups across Present and Absent conditions. Error bars indicate the standard deviation (SD)

### Baseline control measures

In the Present condition, the groups did not differ in baseline activation levels, as measured by STAI subscale for state anxiety [22] (Caffeine group: 35.18 ± 4.69 points, Placebo group: 35.13 ± 5.62 points, *t* (48) = 0.03, *p* = 0.97), nor in their usual daily coffee consumption (Caffeine group: 396.42 ± 205.44 mg, Placebo group: 402.27 ± 177.60 mg, *t* (48) = −0.10, *p* = 0.91), body mass index (Caffeine group: 24.31 ± 3.64 kg/m², Placebo group: 24.72 ± 5.02 kg/m², *t* (48) = −0.33, *p* = 0.73), or time awake at the moment of the Training session (Caffeine group: 7:26 ± 2:48 hours, Placebo group: 6:37 ± 2:41 hours, *t* (37) = 0.93, *p* = 0.35). Additionally, we analyzed whether the groups were comparable in terms of attentional baseline in the Training session (Caffeine group: 7.83 ± 1.57 faces ‘Seen’, Placebo group: 8.05 ± 1.04 faces ‘Seen’, *t* (48) = −0.44, *p* = 0.66). In the Absent condition, the groups did not differ in baseline activation levels, as measured by the state anxiety test (Caffeine group: 36.52 ± 6.50 points, Placebo group: 34.70 ± 7.38 points, *t* (45) = 0.89, *p* = 0.37), nor in their usual daily coffee consumption (Caffeine group: 365.21 ± 201.37 mg, Placebo group: 369.16 ± 217.87 mg, *t* (45) = −0.06, *p* = 0.94), body mass index (Caffeine group: 24.47 ± 3.14 kg/m², Placebo group: 24.38 ± 6.34 kg/m², *t* (45) = 0.06, *p* = 0.95), or time awake at the moment of the Training session (Caffeine group: 7:13 ± 2:41 hours, Placebo group: 6:55 ± 3:16 hours, *t* (39) = 0.32, *p* = 0.74). Additionally, we analyzed whether the groups were comparable in terms of attentional baseline in the Training session (Caffeine group: 8.00 ± 1.82 faces ‘Seen’, Placebo group: 8.54 ± 1.22 faces ‘Seen’, *t* (45) = −1.16, *p* = 0.25).

## Discussion

Contrary to our expectations, caffeine appeared to have a detrimental effect on the consolidation of face recognition memory. This was evidenced in the integrated analysis across Present and Absent conditions, which revealed that the Caffeine group after learning exhibited a significantly lower Correct choice rate and a higher False alarm rate compared to the Placebo group. These effects were consistent across both conditions, indicating a general disruption in recognition accuracy rather than a condition-specific effect. Taken together, the results suggest that caffeine, when administered post-encoding, impairs the ability to reliably discriminate previously seen faces from similar distractors, disrupting rather than enhancing the consolidation of face memory representations. In line with these findings, the CAC curves revealed that the performance of the Caffeine group was consistently inferior to that of the Placebo group across all confidence levels. However, the typical confidence-precision relationship -characterized by increasing accuracy with higher confidence-remained preserved in both groups.

It is important to note that the detrimental effect on face recognition became statistically evident only when both conditions were analyzed together. When assessed separately using a sensitivity measure (*d’*), which accounts for both correct responses (Hits or Correct rejections according to the condition) and false alarms, Caffeine group showed lower than the Placebo group, although this difference did not reach statistical significance.

While previous studies on the effect of caffeine on consolidation have yielded mixed results, to our knowledge, no research to date has reported a significant increase in false alarms in recognition tasks following post-encoding caffeine administration. One potential explanation involves heightened physiological arousal induced by caffeine, which is known to increase the sympathetic nervous system and the HPA axis [23,24,25]. This elevated arousal may enhance the encoding of post-learning stimuli (e.g., faces encountered after the experimental session), increasing interference with consolidation of the original encoded faces. This interpretation aligns with the idea that arousal can both facilitate memory consolidation and increase competition between newly encoded and previously learned information, leading to greater susceptibility to interference [26]. This explanation is consistent with Schwabe et al. (2010) [27], who proposed that arousal not only influences how much we remember but also the quality of what is remembered. Specifically, arousal can shift memory processing from flexible, hippocampus-dependent encoding to more rigid, habit-based learning. If caffeine-induced arousal pushed participants to this kind of processing, it may have reinforced the encoding the new faces encountered after the experiment, at the expense of consolidating the original ones. In our study, the Caffeine group may have unintentionally encoded post-experiment faces more strongly, which interfered with the consolidation of the previously learned faces. This would also be in line with evidence suggesting that caffeine benefits the encoding of incidental information [2].

To our knowledge, the only study that observed a beneficial effect of caffeine on recognition memory consolidation in humans was Borota et al. (2014) [13], which observed that a dose of 200 mg of caffeine could benefit the ability to discriminate between similar and previously seen objects. One possible explanation for the differences between their results and those found in our study could be the substantial difference between the information learned in both studies. In the cited study the stimuli consisted of cartoon-style drawings of objects. These stimuli are rarely found in post-laboratory experiences, that is, they are difficult to compete with stimuli external to the experiment. However, a later replication by Aust & Stahl (2020) [28] found that the enhancing effect reported by Borota et al., (2014) [13] may not be as robust. They proposed that observed improvement could reflect a reversal of the withdrawal rather than true cognitive enhancement. This phenomenon occurs frequently because, by orders from the experimenter, the participants avoid consuming caffeinated beverages before the experiment.

Another possible explanation stems from caffeine’s documented enhancement of global processes [29], which biases individuals towards holistic rather than detail-focused processing. In a recognition memory task, this bias can lead to increased false alarms, as participants may rely on general face similarities rather than discriminating specific features that distinguish a target from a lure. This occurs because global processing could favor the recognition of broad or contextual characteristics of the stimuli instead of discriminating between unique or specific details that differentiate a target stimulus from a lure one.

The effects of caffeine on memory consolidation may also depend on the nature of the stimuli. While object recognition might benefit from enhanced pattern separation mechanisms supported by norepinephrine or long-term potentiation [13,30] face recognition engages a combination of feature and holistic processes [31]. Some evidence suggests that caffeine can elevate hippocampal acetylcholine levels via adenosine A1 receptors [32], potentially disrupting memory consolidation by interfering with the replay of newly acquired memories. This may disproportionately affect face recognition, which is highly sensitive to subtle interference and less reliant on discrete features. If feature traces are weakened during consolidation, participants may rely more heavily on holistic retrieval strategies, increasing the chance of confusing similar faces. This interpretation aligns with findings showing that face recognition accuracy declines when holistic information is disrupted [33]. Thus, caffeine may impair not only the strength of memory traces, but also the cognitive strategies used during retrieval.

Within the limitations of the present work, we did not include an active control group, as recommended by Aust & Stahl (2020) [28], which impeded us from controlling the subjects’ awareness of the pill received. Additionally, we did not measure reaction times, which could have provided further insights into the processing dynamics of the task [34]. Furthermore, it has been observed that impulsivity levels could modulate the effects of caffeine on memory tasks [35] and this variable was not taken into account.

In sum, the findings of this study suggest that caffeine could have a detrimental effect on the consolidation of face recognition memory. However, this effect could be influenced by the unique characteristics of faces as stimuli, given that humans possess a remarkable ability to recognize, encode, and consolidate face information. The facility with which face stimuli are encoded and their constant availability in everyday environments could play a key role in how caffeine modulates

their consolidation. Future research should continue exploring both factors together, as caffeine is the most widely consumed psychoactive substance, and face recognition ability could be crucial in contexts such as the judicial system, where it could aid in the identification of criminals or help avoid wrongful convictions. The present study contributes to understanding how a widely used substance could influence a fundamental human ability.

## Materials and Methods

### Study participants

The study sample consisted of 104 healthy adults between 18 and 40 years old (mean age = 27.30 ± 6.33 years), all of whom were habitual consumers of caffeine beverages. Participants were recruited through advertisements posted on the laboratory social media platforms. Two individuals were excluded from the initial sample for not attending the second session, resulting in a final sample size of 102 participants. Inclusion criteria were: age between 18 and 40 years and no current use of psychotropic medication or history of psychological illness. Exclusion criteria included a history of migraines or chronic headaches, ulcers or gastrointestinal disorders, high blood pressure, or cardiovascular problems. All participants provided written informed consent prior to participation. The study was approved by the Human Ethics Committee (CEH), Faculty of Medicine, University of Buenos Aires (UBA). Participants were instructed to stop consuming caffeine from the night before starting the study and were instructed not to take naps during the two study days.

### Groups

Participants were randomly assigned to one of two groups (Caffeine or Placebo), and to one of two testing conditions (Present or Absent condition).

#### Caffeine group

On Day 1, participants completed the Training session, followed by the administration of 200 mg of caffeine. On Day 2, they performed the Testing session under one of two conditions: Present condition (*N* = 28, 19 females) and Absent condition (*N* = 24, 16 females).

#### Placebo group

On Day 1, participants completed the Training session, followed by the administration of an inert capsule. On Day 2, they performed the Testing session under one of two conditions: Present condition (*N* = 24, 14 females) and Absent condition (*N* = 26, 20 females)

The sample size was determined based on a previous study that examined the effect of caffeine on memory consolidation for a recognition task [13].

### Experimental procedure

The experiments took place in a quiet room using a personal computer. Each participant wore headphones and sat in front of a 24-inch monitor. Before the study, medical staff conducted a health assessment to determine eligibility and the participants completed the questionnaires. Participants then returned to the laboratory on a different day to begin the experiment. On day 1 upon arrival, participants provided written informed consent before beginning the experiment. Then completed the Training session and immediately after received either 200 mg of caffeine or a placebo, administered with a glass of water. On Day 2, at the same time as the previous session, participants returned to the lab to complete the Testing session under identical conditions. All experiments were performed between 10 a.m. and 5 p.m.

### Questionnaires

During the medical interview the BMI (Body mass index) of the subjects was recorded and they were asked about their usual caffeine consumption. To account for caffeine consumed through the local drink mate, we assumed a concentration of 250 mg per liter [36,37], and a value of 200 mg of caffeine per cup of coffee (approximately 240 ml) was assumed [38] Further, although the time of the study sessions was controlled, the specific time of awakening at the time of the Training session was not initially recorded due to an omission and was collected retrospectively. Since this question required retrospective recall, 15 participants were unable to provide an accurate response, so those data were not taken into account for the analysis. Further, prior to the Training session, participants completed the STAI subscale for state anxiety [22] to assess the basal activation state.

### Tasks

During the Training session (Day 1), participants received the following instruction: ‘You will now see a series of faces. The images will automatically scroll. Click the “Start” button and observe them carefully.’ They were then presented with 10 images of human faces (5 males and 5 females, selected randomly), each displayed at the center of the screen for three seconds. Participants were informed that they would later see a series of images and were instructed to classify each as ‘Seen’, ‘Unseen,’ or ‘Familiar.’ During this phase, the same set of 10 images was shown again in random order, and participants were required to make a classification for each one. These response options were included to ensure sustained attention throughout the Training session. Immediately after finishing the Training session, a capsule was administered that could be 200 mg of caffeine or a placebo and the participants left the laboratory. On the following day (Day 2), during the Testing session, participants received the instruction: ‘You are now going to see several groups of faces, each containing 6 photos. Each group could or could not contain one of the faces you watched before. If you recognize any of the faces, select the corresponding item (Each face in the lineup has a visible number ranging from 1 to 6). If you do not recognize any of the faces, select the item ‘I do not recognize any of the photos’. Participants assigned to the Present condition saw a lineup containing the original face along with five similar faces, whereas those in the Absent condition were presented with six faces similar to the original, without the original face included. There was no time limit for the Testing session elections. After making each choice, participants were asked about their level of confidence through the question: ‘How confident are you in your choice? Answer with a number within 1 and 100’.

### Stimuli

On Day 1 (Training session), 10 human face images were generated using Generative Adversarial Networks (GANs) [39]. To create the testing lineups for Day 2 (Testing session), six new faces were generated for each lineup, and transitional images were created between the original face and each of these new faces. From these transitions, we selected the image representing a 50% blend of the original face and a new face, forming the final testing lineups. In the Present condition, the lineup included five transition-generated faces along with the original face from Day 1. In the Absent condition, only the six transition-generated faces were presented. Face image generation and manipulation were performed using a GAN-based architecture, specifically StyleGAN [40], a variant optimized for the synthesis of realistic and controllable images. The implementation was integrated into an interactive web-based platform, enabling the generation of faces with modifiable attributes. This tool allowed for the creation of visual stimuli with controlled variability, ensuring consistency in face structure while exploring different phenotypic characteristics.

### Statistical analysis

Statistical analysis was performed using the IBM SPSS Statistics 25 software. To assess participants’ sensitivity, we used *d-prime* (*d’*), a measure derived from Signal Detection Theory [41]. Unlike raw accuracy, *d’* accounts for both hits (or correct rejections in the Absent condition) and false alarms, providing a bias-free estimate of the ability to discriminate between target and non-target stimuli. In the case of the Present condition, the formula *d*′= Z (Hit Rate) − Z (False Alarm Rate) was used and in the case of the Absent condition *d*′= Z (Correct rejections) – Z (False Alarm Rate) was used. Further, an analysis of variance two-way ANOVA was conducted to assess the effect of caffeine (Caffeine vs. Placebo) and target condition (Present vs. Absent) on correct responses and false alarms (hits/correct rejections). Further, to evaluate the relationship between the accuracy of the elections and the confidence attributed to them, CAC curves were performed. To calculate the value of the correct proportion corresponding to each confidence level (low 0–50%, medium 60–80%, or high 90–100%) the following formula was used: # Correct identifications/# Correct identifications + # Incorrect identifications [42]. Finally, the levels of state anxiety, habitual daily caffeine consumption, body mass index, time awake at the beginning of the training session, and attentional performance (analyzed by comparing the number of face images identified as ‘Seen’ during encoding between groups) were analyzed using Student’s t-tests for both conditions.

## Data availability

The raw data supporting the conclusions of this article will be available after acceptance. The stimuli used are available upon request to the corresponding author.

## Author contribution

C.L. and C.F. made substantial contributions to the conception and design of the work. C.L., A.L.LC and R.J. acquired the data. A.L.LC and R.G. were responsible for the medical screening of participants and administration of the experimental substance. C.L. performed the statistical analyses. C.L. made the graph art. C.L. and C.F. contributed by drafting the work. C.L. and C.F. contributed to revising it critically. C.F. was in charge of funding acquisition, project administration and supervision. All authors contributed to the article and approved the submitted version.

